# Physioxia stimulates extra-cellular matrix deposition and increases mechanical properties of human chondrocyte-derived tissue-engineered cartilage

**DOI:** 10.1101/2020.08.05.229294

**Authors:** J E Dennis, G A Whitney, J Rai, R J Fernandes, T J Kean

## Abstract

Cartilage tissue has been recalcitrant to tissue engineering approaches. In this study, human chondrocytes were formed into self-assembled cartilage sheets, cultured in physiologic (5%) and atmospheric (20%) oxygen conditions and underwent biochemical, histological and biomechanical analysis at one- and two-months. The results indicated that sheets formed at physiological oxygen tension were thicker, contained greater amounts of glycosaminoglycans (GAGs) and type II collagen, and had greater compressive and tensile properties than those cultured in atmospheric oxygen. In all cases, cartilage sheets stained throughout for extracellular matrix components. Type II-IX-XI collagen heteropolymer formed in the neo-cartilage and fibrils were stabilized by trivalent pyridinoline cross-links. Collagen cross-links were not significantly affected by oxygen tension but increased with time in culture. Physiological oxygen tension and longer culture periods both served to increase extracellular matrix components. The foremost correlation was found between compressive stiffness and the GAG to collagen ratio.

**Summary:** Tissue-engineered cartilage formed from human articular chondrocytes produces thicker, stiffer, more extracellular-matrix rich cartilage tissue when grown under physiological (5%) vs. atmospheric oxygen (20%) tension.

## Introduction

Cartilage tissue has very poor intrinsic repair capacity. Whilst osteoarthritis is a complex, multifaceted disease, cartilage degradation is a core component. Autologous chondrocyte implantation and matrix assisted autologous chondrocyte implantation have provided relief to patients but commonly result in fibrocartilage repair^[1]^. Tissue engineering could potentially address this through *in vitro* culture methods to produce functional hyaline cartilage tissue, with several examples currently in clinical trials^[2]^. We, and others, have investigated media supplements and growth factors to improve the expansion and re-differentiation of the expanded chondrocytes^[3–6]^.

It is increasingly apparent that physiological oxygen tension should be the standard culture method to grow tissue engineered human articular cartilage whether it be from mesenchymal stem cells^[7]^, chondrocytes^[8–17]^ or chondroprogenitors^[18]^. The selection of cell type for engineered tissue raises some interesting issues, mesenchymal stem cells commonly progress to hypertrophy^[19]^ as do iPSCs driven down the mesenchymal pathway^[20]^, a negative scenario for the production of hyaline cartilage. Hypertrophy has been reduced but not eliminated during MSC culture^[21]^ and subcutaneous implants by pharmacological and/or culture-dependent methods^[22]^. This study focuses on the use of human articular chondrocytes derived from discarded total joint replacement tissue as both a clinically relevant and non-hypertrophic cell source. We, and others, have focused on scaffold-free self-assembly of tissue-engineered cartilage, as significant similarities to native tissue structure can be achieved^[17, 18, 23–29]^. Significant expansion, up to 8 population doublings, of human chondrocytes whilst maintaining their differentiation capacity has been achieved through their culture on devitalized synoviocyte matrix^[13]^. Using these methods, we produced sheets of human articular chondrocyte-derived cartilage and investigated the effect of low, physiological, oxygen tension and duration of culture on cartilage quality in terms of biomechanics and biochemical content. It was hypothesized that physiological oxygen tension and increased culture duration would improve the mechanical properties of the tissue engineered cartilage through an increase in extracellular matrix content. This study adds to the current body of literature by expanding the donor pool, focuses on chondrocytes isolated from total joint replacement tissue as a cell source, and includes analysis of cartilage-typic collagen heteropolymer formation, collagen cross-links and tensile properties of tissue engineered cartilage.

## Methods

### Cell and tissue culture

Human articular chondrocytes were thawed from frozen stocks obtained from discarded surgical tissue of patients (n=6) undergoing total joint replacement collected with IRB approval. Human articular chondrocytes were expanded under physiological oxygen tension (5%; Physioxia) on synoviocyte derived extracellular matrix in growth media (DMEM-LG supplemented with 10% FBS and 1% penicillin/streptomycin)^[13]^. At the end of first passage (6 experiments) or second passage (2 experiments), cells were trypsinized and seeded at high density (4.4×10^6^ cells/cm^2^) in a custom stainless steel biochamber (Fig. 1)^[28]^. Biochambers were assembled and sterilized by autoclave. The polyester membrane was coated with fibronectin (8 μg/cm^2^ in PBS) and allowed to dry in a biosafety cabinet. Biochambers were used much as previously described but with a stacked, 1.1 cm^2^ circular seeding chamber with longer screws creating greater space above and below the chamber for media exchange (Fig. 1). Biochambers were cultured at either atmospheric oxygen tension (20%; Atm O_2_) or physioxia (5% O_2_) in defined chondrogenic medium (DMEM-HG containing ITS+premix, dexamethasone (100 nM), ascorbate-2-phosphate (120 μM), MEM NEAA, Pen/Strep, glutamax, TGFβ1 10 ng/ml) for 21 days (3-weeks) and 46-56 days (7-weeks) with 50% media changes every other day. All cultures were grown at 37 °C with 5% CO_2_ in a humidified atmosphere. After 5-days in static culture, all biochambers were put on a rotating shaker (60 RPM). At the end of culture, 3 five mm skin biopsy punches were taken for mechanical assessment (equilibrium modulus and tensile modulus). Remaining tissue was assessed by biochemical (glycosaminoglycan (GAG), DNA, hydroxyproline (HDP), and collagen cross-link content (hydroxylysyl pyridinoline + lysyl pyridinoline; HP+LP; moles/mole collagen)) and histological assays. *Biochemical assays:* GAG, DNA and HDP assays were conducted much as previously described^[17]^. Briefly, tissue engineered cartilage was digested with papain solution (25 μg/mL papain, 2 mM cysteine, 50 mM sodium phosphate, and 2 mM EDTA adjusted to pH 6.5 [all from Sigma-Aldrich]) at 65 °C for 3h. Digested tissue was then split between the HDP assay and GAG/DNA assays. For the GAG/DNA portion of the assay, papain digested samples were inactivated by sodium hydroxide (2 volumes 0.1 M NaOH) then neutralized with acidified phosphate buffer (2 volumes 100 mM sodium phosphate pH 7.2 acidified with 0.1 M HCl). GAG was assessed using safranin-O: Neutralized samples were incubated with Safranin-O solution (0.05% in 50 mM sodium acetate) on a dot blot apparatus (BioRad) with a 0.45 μm nitrocellulose membrane in duplicate. Dots were punched from the membrane and incubated in cetylpyridinium chloride solution (10%, Alpha Aesar) at 37 °C for 20 min. The extract was then transferred in triplicate to a clear 96-well microplate and absorbance measured (536 nm; Tecan M200). GAG concentrations were calculated from a standard curve produced with chondroitin sulphate (Seikagaku Chemicals). DNA was assessed from neutralized samples using buffered Hoechst solution (0.667 μg/ml in 0.2 M pH 8.0 phosphate buffer; 33258; Sigma-Aldrich). Neutralized digest (20 μl) was transferred to a black 96-well plate in duplicate and Hoechst solution added (100 μl) then fluorescence read (Ex 365 nm, Em 460 nm; Tecan M200). DNA content was calculated from a standard curve (calf thymus DNA; Sigma-Aldrich) made in neutralized papain buffer. For the assessment of HDP, papain digested samples were acid hydrolyzed overnight (10:1 vol/vol, 6 M HCl, 110 °C). Acid hydrolysate was then evaporated to dryness by incubation at 70°C 1-2 days. Samples and hydroxyproline standards were then resuspended in ddH_2_O, mixed with 1 volume copper sulphate (0.15 M), 1 volume NaOH (2.5 M) and incubated (50°C, 5 min). Samples were then oxidized by incubation with hydrogen peroxide (1 volume, 6% H_2_O_2_; 50°C, 10 min). To this solution, 4 volumes of sulphuric acid were added (1.5 M H_2_SO_4_) then reacted with Ehrlich’s reagent (2 volumes: 10% w/v 4-dimethylamino benzaldehyde in 60% isopropanol, 26% perchloric acid, 14% MΩ water) at 70°C for 16 min. After cooling, samples and standard absorbance was read on a plate reader (505nm, Tecan M200). Collagen content was estimated from hydroxyproline concentration by a conversion factor of 7.6^[30]^.

**Figure 1.**
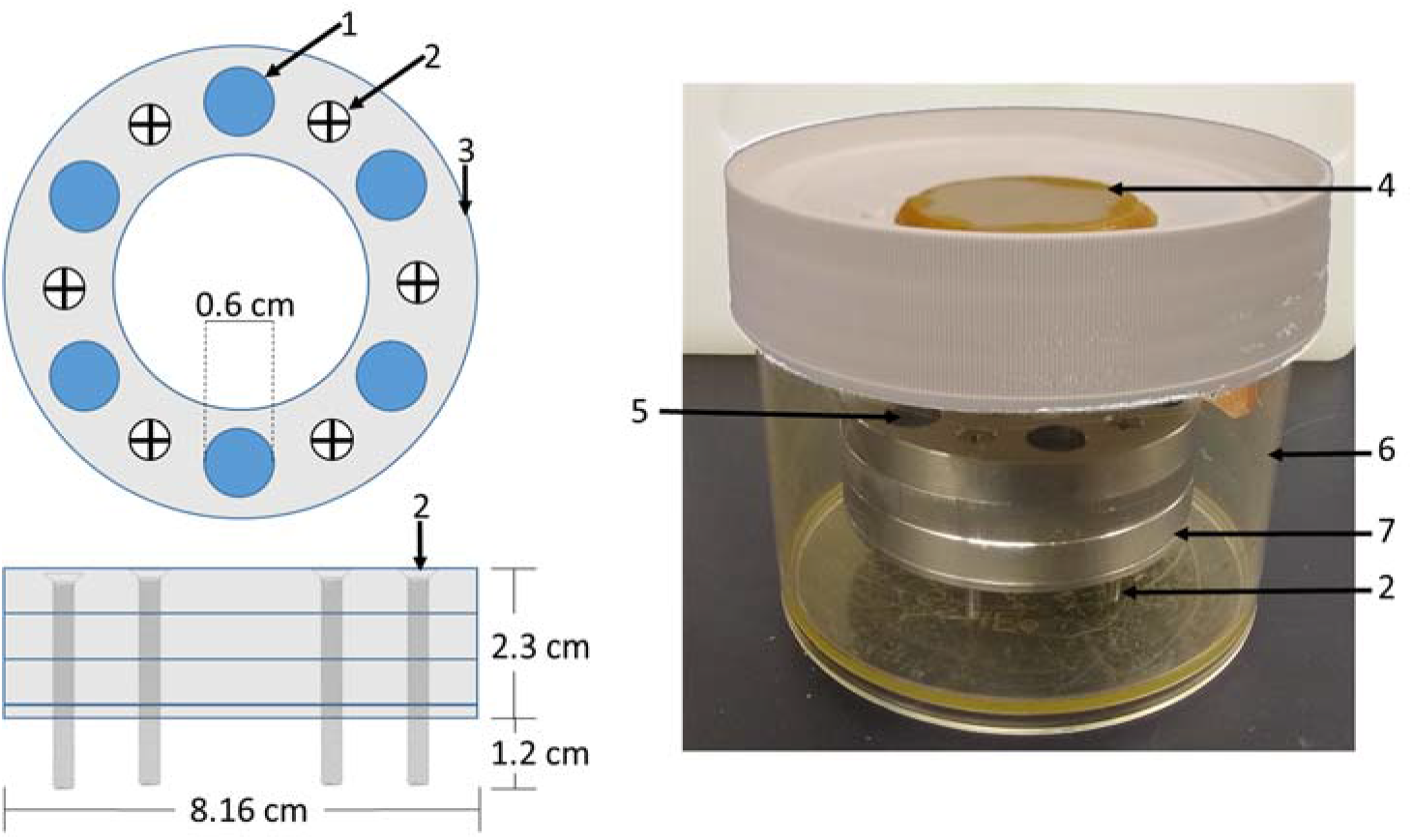
Biochamber model and setup. A circular biochamber design with a 1.13 cm^2^ cell seeding area (1) was used with three seeding chambers (3) stacked on top of each other giving a 2.1 ml seeding volume. Stainless steel screws (2) were used to raise the biochambers 1.2 cm allowing media access to the cell sheet through the polyester membrane (7) sandwiched between the bottom 0.2 cm plate and the seeding chambers. Biochambers were assembled and placed in Nalgene containers (6) with a ceramic filter on the lid (4). Defined chondrogenic media was added to the level of the membrane, cells seeded, then media added to 0.2 cm below the top of the topmost seeding chamber.

#### Collagen cross-link analysis

Samples were dried, weighed and then acid hydrolyzed in 6M HCl, 110 °C for 24h. Hydroxylysyl pyridinoline (HP) and Lysyl pyridinoline (LP) cross-linking residues were resolved and quantified by C-18 reverse phase HPLC with fluorescence detection (excitation 297 nm, emission 396 nm) and total collagen content was determined as described^[31]^.

#### Collagen heteropolymer analysis

Unused portions of the samples used for mechanical analyses were used to qualitatively fingerprint cross-linked collagen types by western blots. The heteropolymeric collagen network formed in the samples was depolymerized in equal volumes of 0.5 M acetic acid containing 100 μg/ml pepsin for 18 h at 4 °C. Equal aliquots of solubilized collagen were analyzed by SDS-PAGE and the separated collagen chains visualized by Coomassie blue staining. Pepsin-extracted type II collagen from rabbit articular cartilage was used as a control. The separated collagen chains were also blotted onto PVDF membranes and probed with mAb 1C10 to identify α1(II) chains and with mAb 10F2, pAb 5890, mAb 2B4 to identify collagen chains cross-linked to the C-telopeptide of α1(II), to the N-telopeptide of α1(XI) chains and to the α1(IX) chains respectively^[32]^. As we have described before, this validates if a heteropolymer of type II and type XI collagen had formed^[33]^.

### Histological analysis

Samples were fixed in neutral buffered formalin overnight at 4°C then switched to PBS at 4°C until embedded. Samples were embedded by sequential dehydration in graded ethanols, xylene and paraffin. Paraffin sections (8 μm) were deparaffinized and hydrated before staining with safranin-O (Sigma-Aldrich) for GAG with a Fast Green (AA16520-06, Alfa Aesar) counterstain. For immunohistochemistry, hydrated sections were subjected to antigen retrieval by pronase (1 mg/ml in PBS containing 5 mM calcium chloride; Sigma-Aldrich) incubation for 10 minutes at room temperature. Primary antibodies against type I collagen (631703, MP Biomedical, 1:1000), type II collagen (DSHB II-II6B3 cell culture supernatant, 1:500) and type X collagen (kind gift of Gary Gibson, Henry Ford Hospital, Detroit, MI; 1:500) were incubated with tissue sections at 4°C overnight. Sections were then rinsed and stained with secondary antibody (biotinylated horse anti-mouse; Gibco; BA2000; 1:2000) for 1h at room temperature before rinsing and incubation with streptavidin-HRP (SNN1004, Invitrogen, 1:5000) 30 min at room temperature. Detection was then made with VIP substrate (Vectashield) by incubation at room temperature for 10 minutes. Slides were rinsed and counterstained with Fast Green before mounting.

### Mechanical analysis

Samples were thawed in PBS solution equilibrated to room temperature for at least 30 minutes. Punches were measured 3 times with digital calipers to assess thickness. Compressive equilibrium moduli were determined as previously described^[34]^. Briefly, after an initial tare load of 0.2 N, 4 sequential strains of 5%, 10%, 15%, and 20% were applied, with a stress-equilibration period of 30 minutes between each strain step. The stress measured at the end of each strain period was taken as the apparent stress at the corresponding strain level. The equilibrium modulus was determined as the slope between the apparent stress and the strain.

To test elastic tensile Young’s moduli, a custom dogbone punch was made from skin biopsy punches and punches taken from the 5mm punch (Fig. 2A). Custom holders were made from overhead projector sheets (Supplemental file 1) and dogbones attached using cyanoacrylate glue (Fig. 2A4), tissue hydration was maintained with PBS. Tissues were stretched to failure (Fig. 2B and 2C) and the tensile Young’s modulus, ultimate tensile stress and yield stress calculated. Residual pieces of cartilage from the dogbone punch were used for collagen typing and heteropolymer analysis.

**Figure 2.**
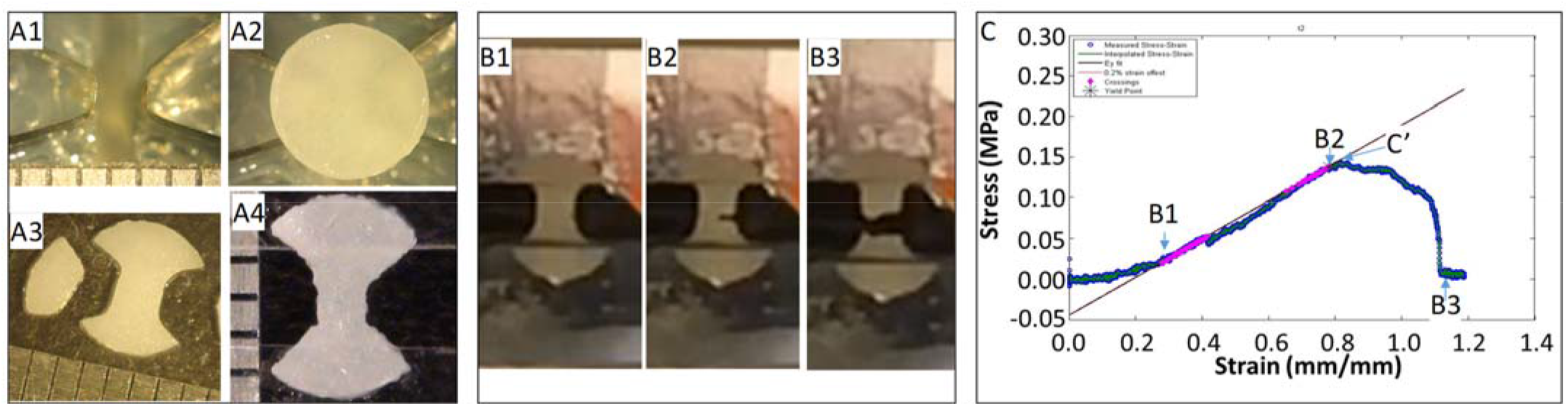
Tissue engineered cartilage sample preparation and tensile testing. Panel A shows the custom manufactured dog bone punch (A1), the 5mm punched tissue-engineered human cartilage sheet on the dogbone punch (A2), the resulting dogbone (A3) and the dogbone fixed to the OHP film adapter to insert into the grips of the mechanical testing device. Panel B shows sequential images of a piece of tissue-engineered human cartilage being tested in tension to failure. Panel C shows the stress strain curve of the cartilage piece with approximate points shown in the graph of the images in B, with B2 indicating the yield stress, in this case close to the ultimate yield stress (C’), and the best fit line showing the tensile Young’s modulus.

### Samples and Statistics

Eight separate experiments were performed with chondrocytes from 6 human donors. For the analysis of GAG/DNA/HDP, 3 samples per sheet were taken from up to 2 sheets and assayed in duplicate. Each experiment was averaged and data shown represents the average for the experiment. When sheets were too flimsy to be manipulated, GAG/DNA/HDP was not performed (3 experiments, 3 donors, 5 data points, all atmospheric conditions). A mixed-effects model was used to analyze log transformed values with Sidak’s multiple comparisons test (GraphPad Prism, V8.4.2). Collagen crosslink density was analyzed using a mixed-effects model with Sidak’s multiple comparisons test (GraphPad Prism, V8.4.2).

For mechanical tests two sheets were made for each donor and 2-3 five mm biopsy punches were taken from each sheet. Punch thickness was measured with digital calipers 3 times and the average value taken for the thickness. Thickness data was analyzed by a mixed-effects model with Sidak’s multiple comparisons test (GraphPad Prism, V8.4.2). Punches that were incomplete circles, curled too much to get a flat sheet in testing were excluded from analysis. Atmospheric oxygen tension sheets from donors that were insufficiently sturdy or were too thin to be manipulated (2 of 6) have the imputed compressive modulus value of the weakest sheet tested (0.3129 kPa). All data represent the average of the samples tested (n = 1-4) for each independent experiment (n = 8). Data were analyzed by repeated measures two-way ANOVA with Sidak’s multiple comparisons test (GraphPad Prism, V8.4.2). Four donors were included in the tensile testing experiments, 2 of which failed to form sheets that were sturdy enough to be tested when cultured at atmospheric oxygen tension. Data were analyzed by two-way ANOVA with Sidak’s multiple comparisons test (GraphPad Prism, V8.4.2).

## Results

Handleable tissue-engineered human cartilage sheets formed in all 8 experiments at physiological oxygen tension vs. 5 of 8 at week 3 and 6 of 8 by week 7 at atm O_2_.

### Biochemical assays

There was a significant increase in extracellular matrix content, both in terms of GAG/DNA (Fig. 3A) and collagen/DNA in human tissue-engineered cartilage sheets when cultured in physioxia vs. atm O_2_ (Fig. 3B). This increase in GAG/DNA was not significantly affected by time in culture (Fig. 3A). Only under physioxia was the accumulation of collagen/DNA greater at the 7-week time point (Fig. 3B).

**Figure 3.**
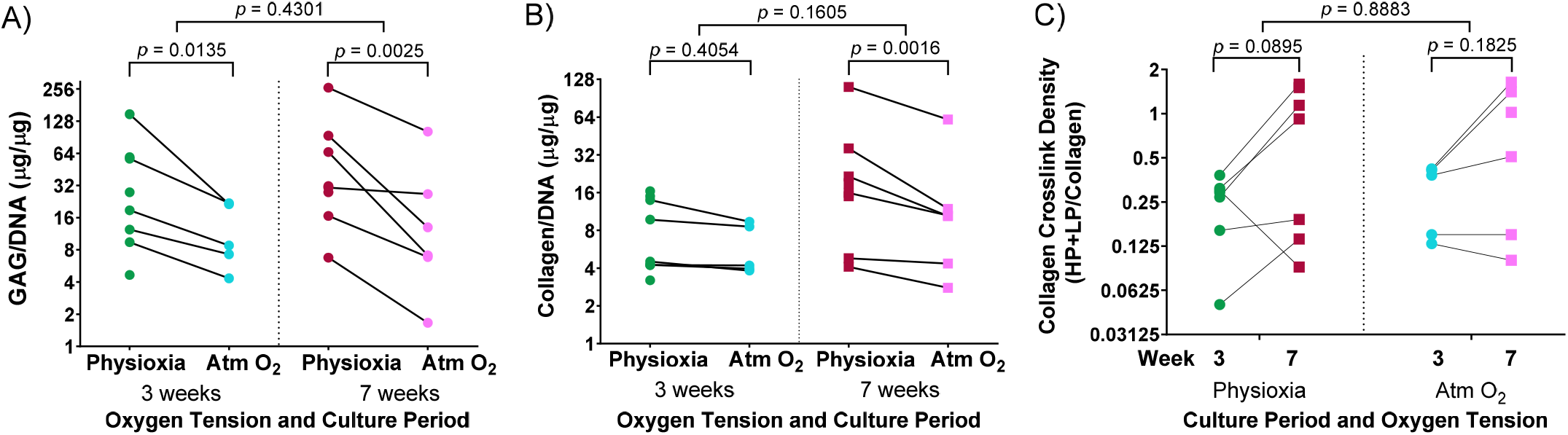
Extracellular matrix deposition in tissue-engineered human cartilage sheets. Tissue-engineered human cartilage sheets had higher glycosaminoglycan content in Physioxia vs. Atm O_2_, time in culture had no appreciable effect (A). Sheets had more collagen when grown in Physioxia at 7 weeks (B). Trivalent Collagen collagen cross-links density increased over time, with no appreciable effect of oxygen tension (C). Note, all measures are shown on a log (base 2) scale.

Collagen trivalent hydroxylysyl and lysyl pyridinoline cross-links were formed in culture at both oxygen tensions and increased with time in culture, with no apparent effect of oxygen tension (Fig. 3C). To determine if cartilage-specific collagen type II-IX-XI heteropolymer formed in culture we used specific antibodies to qualitatively fingerprint cross-linked collagen chains as we previously established^[33]^. Fig. 4A. shows an SDS-PAGE gel of pepsin extracted collagen from human tissue engineered cartilage grown under Atm O_2_ and Physioxia. Purified human type II collagen (lane1) and tissue engineered human cartilage is shown in the lanes 2-5. Lanes 2, 4 and lanes 3, 5 were normalized to wet weight of unused portion of neo-cartilage after mechanical tests. Only a qualitative evaluation of collagen in these samples was possible. The major Coomassie blue stained pepsin-resistant chain observed was the α1(II) collagen chain which migrates similarly to the chain seen for human type II collagen purified from adult articular cartilage. Two other pepsin-resistant chains of varying intensities were observed (lanes 2-5), migrating above the α1(II) chains. The chains migrate similarly to the α1(XI) and α2(XI) chains in type XI collagens. The α2(I) and β2(I) chain characteristic of type I collagen is a minor component of the tissue engineered cartilages.

**Figure 4.**
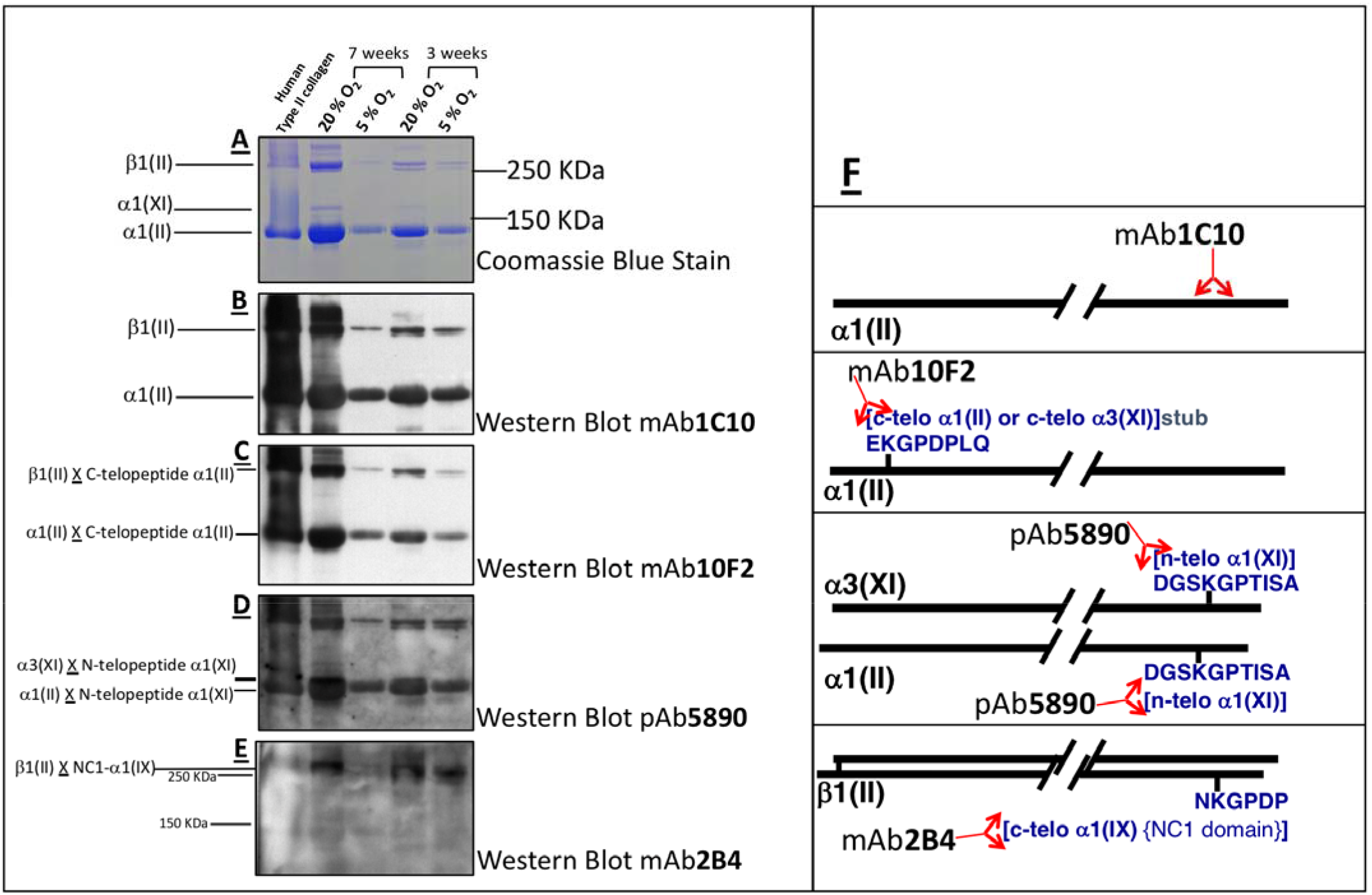
Collagen heteropolymer crosslink analysis. Collagen in human tissue-engineered cartilage was analyzed by SDS page and western blot. Lane 1) human type II collagen (control); 2) 7-week Atm O_2_; 3) 7-week Physioxia; 4) 3-week Atm O_2_; 5) 3-week Physioxia. Panels: A) Coomassie blue stain; B) Western blot type using II collagen antibody to α1(II) chain in helical region; C) Western blot using antibody to C-telopeptide of α1(II) chains of type II collagen; D) Western blot using antibody to N-telopeptide of α1(XI) chains of type IX collagen; E) Western blot of antibody that recognizes C-telopeptide of the non-collagenous domain of α1(IX) chains of type IX collagen of type IX collagen. F) Molecular interpretations of collagen heteropolymer assembly from Western blot analysis. Locations of the epitopes in the α1(II) chain or in telopeptide stubs cross-linked to the chains are also shown

Western Blot using the antibody 1C10 (Fig 4B) confirmed the α1(II) and the β1(II) chains of type II collagen (lanes 2-5) indicating the chondrocytes elaborated an extensive extracellular matrix containing type II collagen. The antibody 10F2 reacted with the α1(II) chain as expected for a cross-linked type II collagen polymer (Fig 4C), (C-telopeptide of type II collagen cross-linked to α1(II) collagen chains) indicating a crosslinked collagen network assembled in the all the neo-cartilages. Following an extended exposure, the antibody also reacted with α1(XI) collagen chains implying this chain was cross-linked to the C-telopeptide of type II collagen and that type XI collagen was copolymerized and cross-linked to C-telopeptides of type II collagen (data not shown). A similar blot when probed with the antibody 5890 clearly reacted with the α1(II) chain and the α3(XI) chain in the neo-cartilages (Fig 4D). (The α1(II) and α3(XI) chains are the identical product of the type II collagen gene but post translational modifications causes the chains to migrate differently on SDS-PAGE). This indicated that the N-telopeptide of the α1(XI) collagen chain was cross-linked to the α3(XI) and the α1(II) chain and thus a heteropolymer of type XI-type II collagen molecules had formed. The antibody 2B4 strongly reacted with the β1(II) chain of type II collagen in the neo-cartilages (Fig 4E) indicating that this chain was cross-linked to the α1(IX) chain of type IX collagen and a heteroplymer of type IX-type II had formed. This fingerprint pattern on western blots showed that a cross-linked heteropolymer of type II-IX-XI had assembled in the neo-cartilages. Fig 4F shows molecular interpretations of collagen heteropolymer assembly from Western blot analysis. Locations of the epitopes in the α1(II) chain or in telopeptide stubs cross-linked to the chains are also shown^[35]^.

### Mechanical Assays

Tissue-engineered human cartilage sheet thickness increased with time in culture at both physiological and atmospheric oxygen tensions (Fig. 5A). Sheets produced in physioxia were significantly thicker than those produced in atm O_2_ (Fig. 5A). Compressive stiffness of the cartilage sheets was greater in sheets grown in physioxia at both 3-weeks and 7-weeks (Fig. 5B), time in culture was only a significant factor for sheets grown in physioxia. Tensile stiffness of sheets increased with time in culture at both physiological and atmospheric oxygen tensions (Fig. 5C), for this measure there was no appreciable effect of oxygen tension. The highest correlation for compressive mechanical stiffness with biochemical measures was achieved with the ratio of GAG/collagen (D). None of the biochemical data showed significant correlation with elastic tensile modulus (Data not shown).

**Figure 5.**
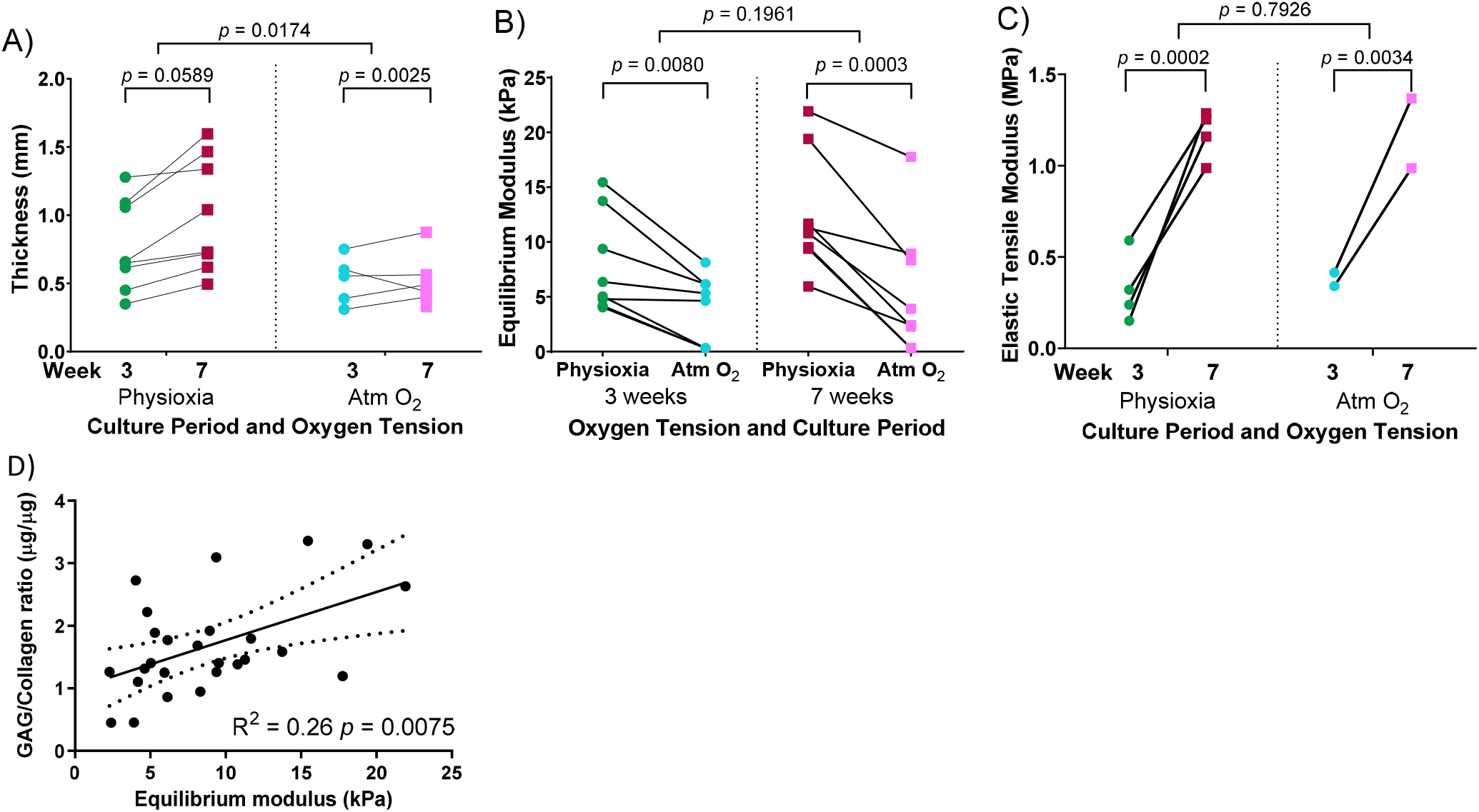
Mechanical testing of tissue-engineered human cartilage sheets. Tissue-engineered human cartilage sheets grew significantly thicker over time in both Physioxia and Atm O_2_, with thicker sheets overall in Physioxia (A). Sheets were stiffer in compression when grown at physiological oxygen tension vs. Atm O_2_ both at 3 weeks and 7 weeks (B). When tested for tensile properties, sheets gained tensile stiffness over time but oxygen tension had no appreciable effect (C). Equilibrium modulus correlated with GAG/Collagen ratio (D).

### Histological analyses

Tissue-engineered human cartilage sheets were stained for glycosaminoglycan content (safranin-O) and collagen types I, II and X. At the 3-week time point, sheets were thicker when grown in physioxic conditions vs. atmospheric oxygen tension (0.77 ± 0.33 vs. 0.52 ± 0.18 mm; mean ± S.D.; Figs. 6 and 5A). Thickness increased over time in physioxic conditions but not in atmospheric oxygen tension (1.0 ± 0.42 vs. 0.52 ± 0.19 mm; mean ± S.D.). Glycosaminoglycan staining was more intense in sheets grown under physioxic conditions (Fig. 6A2) vs. atmospheric oxygen tension (Fig. 6A1). Type I collagen and type II collagen staining was similar under both oxygen tensions at 3-weeks (Fig. 6B1, 6B2, 6C1, 6C2). Type X collagen staining was slightly increased under atmospheric oxygen tension (Fig. 6D1) vs. physioxic (Fig. 6D2) conditions at week 3. At the 7-week time point, safranin-O staining in sheets grown in atmospheric oxygen tension (Fig. 6A3) had increased to a similar level as sheets grown in physioxic conditions (Fig. 6A4). Type I collagen staining under atmospheric oxygen tension at week 7 (Fig. 6B3) has decreased intensity vs the 3-week time point (Fig. 6B1). Similarly, the sheets grown under physioxic conditions at week 7 have reduced or minimal staining for type I collagen (Fig. 6B4) vs. (Fig. 6B2). Type II collagen was relatively intense under both oxygen tensions (Fig. 6C3 and 6C4). Type X collagen staining was more intense on the upper surface of the sheet grown under atmospheric oxygen tension at week 7 (Fig. 6D3). In sheets grown under physioxic conditions, type X collagen staining was predominantly intracellular vs. extracellular (Fig. 6D4).

**Figure 6.**
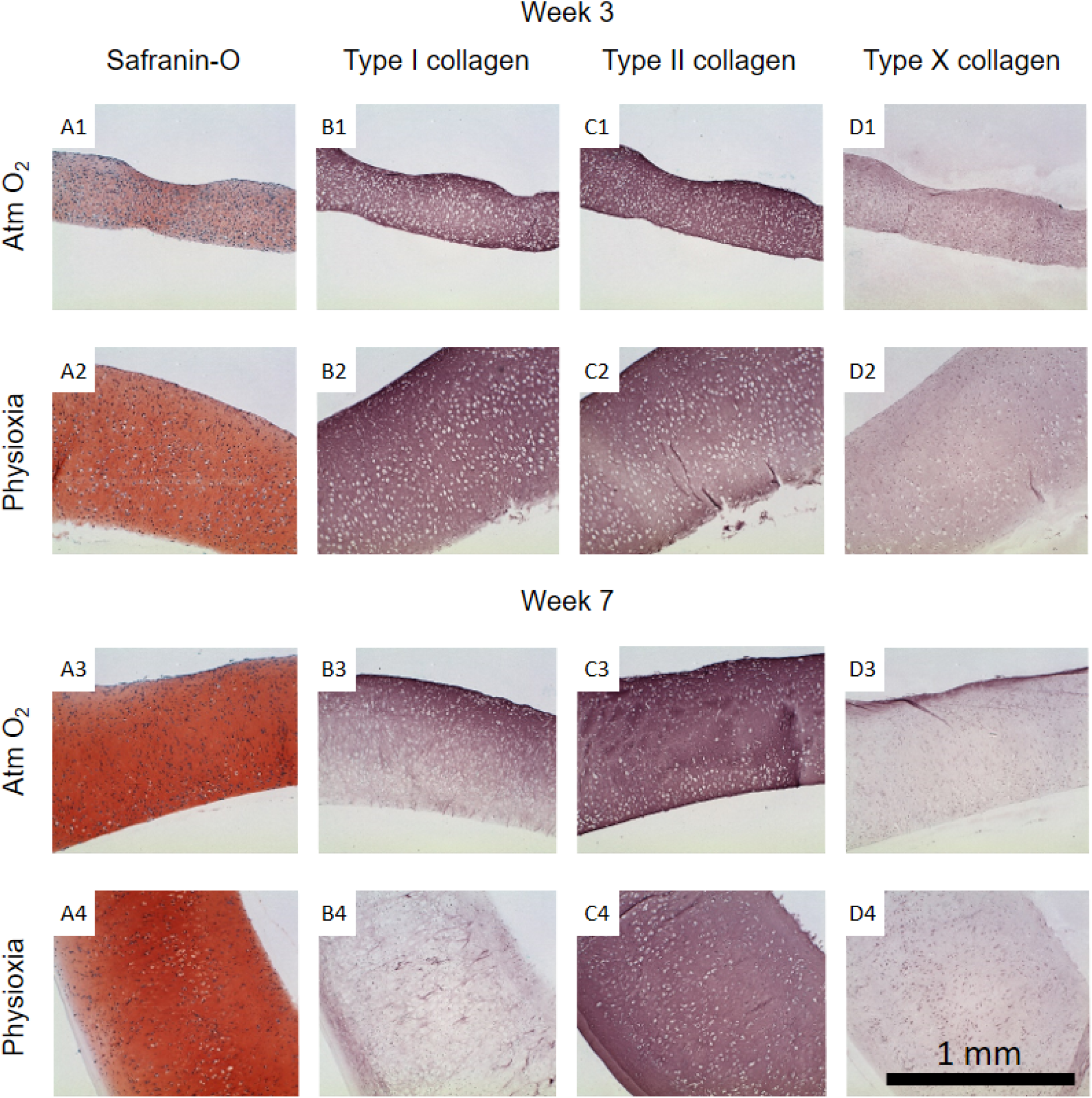
Histological analysis of human tissue-engineered cartilage sheets. Tissue-engineered human cartilage sheets from a single donor are shown (see supplemental data for other donors), they were stained for glycosaminoglycan (safranin-O, column A), type I collagen (column B), type II collagen (column C) and type X collagen (column D). Rows 1 and 2 show data from the 3-week time point and rows 3 and 4 show data from week-7. Atmospheric O_2_ sheets are shown in rows 1 and 3, physioxic sheets are shown in rows 2 and 4. The scale bar indicates a 1 mm distance.

## Discussion

Critically, scaffold-free human tissue-engineered cartilage sheets were successfully formed in physioxic conditions for all donors. There was a large degree of variation between donors in terms of glycosaminoglycan deposition, collagen deposition and crosslink density. Even with this wide variation in donor response, a consistent effect of increased glycosaminoglycan deposition through growth under physioxic conditions was remarkable. Similarly, a consistent increase in total collagen deposition was also found through culture under physiological oxygen conditions. Unfortunately, the biochemical assay for collagen does not discriminate between the different types of collagen. This shortcoming is apparent when looking at the histological data, where temporal and regional variation in collagen type and intensity are evident. Initial expression and replacement of type I collagen has been documented developmentally *in vivo*^[36, 37]^ and, as we have also shown in human bone marrow derived stem cell neo-cartilage, engineered *in vitro*^[33]^. This could indicate that expression and replacement is a normal progression in tissue-engineered cartilage development and that replacement of type I with type II collagen is aided by physioxia. There was no apparent effect of oxygen tension on trivalent collagen cross-linking, but significant increases in total hydroxylysyl and lysyl pyridinoline cross-link formation were observed with longer culture duration. This indicates that under both physioxic and normoxic conditions a fibrillar network of collagen with mature cross-links had formed in neo-cartilage. Western blot analyses using established antibodies to specific collagen peptides involved in covalent cross-link formation^[32, 35]^ indicated that a cross-linked hetropolymer of type II-IX-XI collagen had formed in the tissue engineered cartilage. This cross-linked collagen heteropolymer is typical of cartilage and is essential in the proper assembly of the cartilage collagen fibril^[38]^. Our findings that a similar nascent heteropolymeric template is formed in human neo-cartilage with increased cross-linking with time in culture point to a progressive formation of type II collagen based fibril network typical of cartilage.

Subjective assessment of the sheets (physical handling) indicate that longer culture durations gave stronger sheets in all cases, although this was not fully supported by the equilibrium moduli which only showed a benefit in sheets grown under physioxia. This is potentially due to untestable sheets formed under atmospheric oxygen culture (11 of 16 sheets). However, the results did show an increase in both total collagen content and collagen cross-link content with time. These increases correlated with the increased tensile properties of the tissues. Makris et al. (2013) also found that hypoxia increased collagen crosslinks in tissue engineered bovine cartilage constructs but that they also found weak correlations to compressive mechanical properties^[39]^. The combined increase in compressive and tensile stiffness, along with the greater accumulation of extracellular matrix components, through culture under physioxia leads us to recommend this culture method for chondrogenic experiments.

There remain significant challenges in producing autologous, tissue-engineered cartilage. Current clinical trials cover a wide range of approaches, many using allogenic cells (for review see ^[2]^). The advantages of a well characterized allogenic cell bank are clear given the range of GAG and collagen contents due to donor variability. In chondrocyte progenitor experiments, this has been further focused in on showing clonal variability^[40]^. Interestingly, while there was a wide range of extracellular matrix component concentrations detected, the distribution in mechanical properties was actually relatively narrow. Evans et al. (2005) found that cartilage compressive stiffness correlated with GAG density in native tissues^[41]^. Roeder et al. found that tensile modulus increased with increasing concentration of type I collagen engineered constructs in a linear manner^[42]^. Our data showed the greatest correlation of biochemical measures with compressive moduli when the GAG/Collagen ratio was used but, probably due to the 4 conditions and 6 donors, this correlation was relatively weak. When looking at biochemical content correlations with tensile properties, nothing gave a significant correlation; this analysis was hampered by the number of conditions analyzed and lack of sheet formation for two of the 4 donors at atmospheric oxygen tensions. Similarly, Williamson et al. (2003) found no significant correlation between bovine fetal cartilage and tensile tests^[43]^. Overall, the data indicate that physiological oxygen tension is beneficial to chondrogenesis of human tissue-engineered cartilage sheets formed through scaffold-free culture of human articular chondrocytes.

## Conclusions

Tissue-engineered human cartilage sheets, formed through scaffold free self-assembly of articular chondrocytes, have significantly more extracellular matrix with correlative increases in compressive stiffness. Temporal increases in collagen crosslinks and type were more evident in sheets grown under physiological oxygen tension and these correlated with increased tensile stiffness.

## Acknowledgements

This work was supported in part by NIH grants AR053622 (J.E.D), DE015322 (J.E.D.), AR057025 (R.J.F.) and AR037318 (J.R. and R.J.F.). The authors would like to thank Geoffrey R. Traeger for his technical expertise and Dr. Gary Gibson (Henry Ford Hospital) for the type X collagen antibody.

